# Immunological factors, but not clinical features, predict visceral leishmaniasis relapse in patients co-infected with HIV

**DOI:** 10.1101/2021.03.30.437646

**Authors:** Yegnasew Takele, Tadele Mulaw, Emebet Adem, Caroline Jayne Shaw, Susanne Ursula Franssen, Rebecca Womersley, Myrsini Kaforou, Graham Philip Taylor, Michael Levin, Ingrid Müller, James Anthony Cotton, Pascale Kropf

## Abstract

Visceral leishmaniasis (VL) has emerged as a clinically important opportunistic infection in HIV patients, as VL/HIV co-infected patients suffer from frequent VL relapse. Here, we followed cohorts of VL patients with or without HIV co-infections in Ethiopia and collected detailed clinical and immunological data during 12 months of follow-up. By the end of the study 78.1% of VL/HIV patients, but none of the VL only patients, had relapsed. Despite clinically defined cure, VL/HIV patients maintained high parasite loads, low BMI, hepatosplenomegaly and pancytopenia throughout follow-up. During detailed immunological study throughout the follow-up period, we identified three markers associated with VL relapse: i) failure to restore antigen-specific production of IFNγ, ii) persistently low CD4^+^ T cell counts, and iii) high expression of PD1 on CD4^+^ T cells. We show that these three markers combine well in predicting VL relapse, and that all three measurements are needed for optimal predictive power. These three immunological markers can be measured in primary hospital settings in Ethiopia and can predict VL relapse after anti-leishmanial therapy. The use of our prediction model has the potential to improve disease management and patient care.

## INTRODUCTION

Visceral leishmaniasis (VL) is one of the most neglected tropical diseases. An estimated 550 million individuals are at risk of VL in high-burden countries and 17,082 new cases of VL were reported in 2018, with Brazil, Ethiopia, India, South Sudan and Sudan, each reported >1000 VL cases, representing 83% of all cases globally (1). These numbers are widely acknowledged to underestimate the real burden because of the remote location of areas endemic for VL and the lack of surveillance. VL inflicts an immense toll on the developing world and impedes economic development, with an estimated annual loss of 2.3 million disability-adjusted life years (2). In Ethiopia, where this study took place, VL is one of the most significant vector-borne diseases; over 3.2 million people are at risk of infection (3). VL is caused by infections with parasites of the *L. donovani* species complex, but the majority of infected individuals control parasite replication and do not progress to disease. Some individuals will progress and develop VL that is characterised by hepatosplenomegaly, fever, anaemia and wasting; this stage of the disease is generally fatal if left untreated (4, 5). Following the HIV-1 pandemic, VL has emerged as an opportunistic infection: VL accelerates the progression of HIV infection to AIDS, and conversely HIV infection increases the risk of developing symptomatic VL (6, 7). Ethiopia has the highest rate of VL/HIV co-infections in Africa, with HIV present in up to 30% of VL cases (8).

HIV co-infection present a major challenge in the prevention and control of VL (9, 10): VL/HIV co-infected patients experience higher rates of treatment failure, drug toxicity, mortality and VL relapse rates compared to patients with VL alone (10, 11). In Ethiopia, over 50% of VL/HIV co-infected patients will experience relapse of VL between 3 and 9 months post anti-leishmania treatment (12). The mechanisms accounting for the increased rate of VL relapse in VL/HIV co-infected patients are poorly characterised. Markers such as low CD4^+^ T cell counts, high parasite loads at the time of VL diagnosis and during follow-up, not being on antiretroviral therapy (ART) at the time of VL diagnosis and *Leishmania* antigenuria have shown variable degrees of prediction accuracy (9, 13–15). Another predictive marker of VL relapse in VL/HIV co-infected patients is a previous history of VL relapse (16).

One of the main immunological characteristics of VL patients is their profound immunosuppression (17): these patients do not respond to the Leishmanin skin test, their peripheral blood mononuclear cells (PBMCs) have an impaired capacity to produce IFN-γ and to proliferate in response to *Leishmania* antigen; this dysfunctional response to antigenic challenge is restored following successful chemotherapy (18) and reviewed in (19–21). The mechanisms leading to impaired T cell responses in VL patients remain to be fully understood.

Our knowledge of the immunopathology of VL/HIV co-infections is particularly sparse. Based on the current literature, it appears that the failure to control parasite replication results in chronic inflammation that leads to exhaustion of the immune system and failure to generate efficient T cell responses. Little is known about the immunological parameters associated with successful therapy. At the end of treatment, the discharge of VL/HIV co-infected patients from hospital is based on clinical and parasitological cure (22). However, no clinical sign predicts increased risk of relapse (22).

Here, we followed two cohorts of VL and VL/HIV co-infected patients in Ethiopia and collected detailed clinical and immunological data during 12 months of follow-up.

Genetic variation between parasites or re-infection of VL/HIV patients could be responsible for the increased rate of relapse. However, genomic data from isolates taken from the same patient cohorts as in the present study shows that infections in VL and VL/HIV patients are caused by parasites from the same population and that almost all relapses are caused by recrudescence of the initial infection, rather than re-infection with a different isolate (23). We thus aim to generate the most detailed picture to-date of the natural history of VL and VL/HIV infections in Ethiopia and to identify clinical and immunological markers associated with VL relapse during follow-up. These markers need to be suitable for measurement in a primary hospital setting in Ethiopia, so that they could contribute to improved evaluation of treatment success and ultimately improve the extremely poor outcomes for these patients.

## RESULTS CLINICAL DATA

### Frequency of VL relapse in VL and VL/HIV patients

We followed VL/HIV patients for up to 3 years and compared their rates of VL relapse with those of VL patients. During this study, three VL/HIV patients left the treatment centre before the end of treatment (EoT) and five died during treatment; 41 were treated successfully, i.e. had a negative test of cure. Following anti-leishmanial treatment, nine were lost to follow-up and 32 VL/HIV patients were followed for up to 3 years, out of these, 25 (78.1%) experienced at least one episode of VL relapse.

This was in sharp contrast with VL patients; four VL patients died during treatment and 10 were lost to follow-up, but none of the successfully treated VL patients experienced VL relapse (Figure 1).

**Figure 1:**
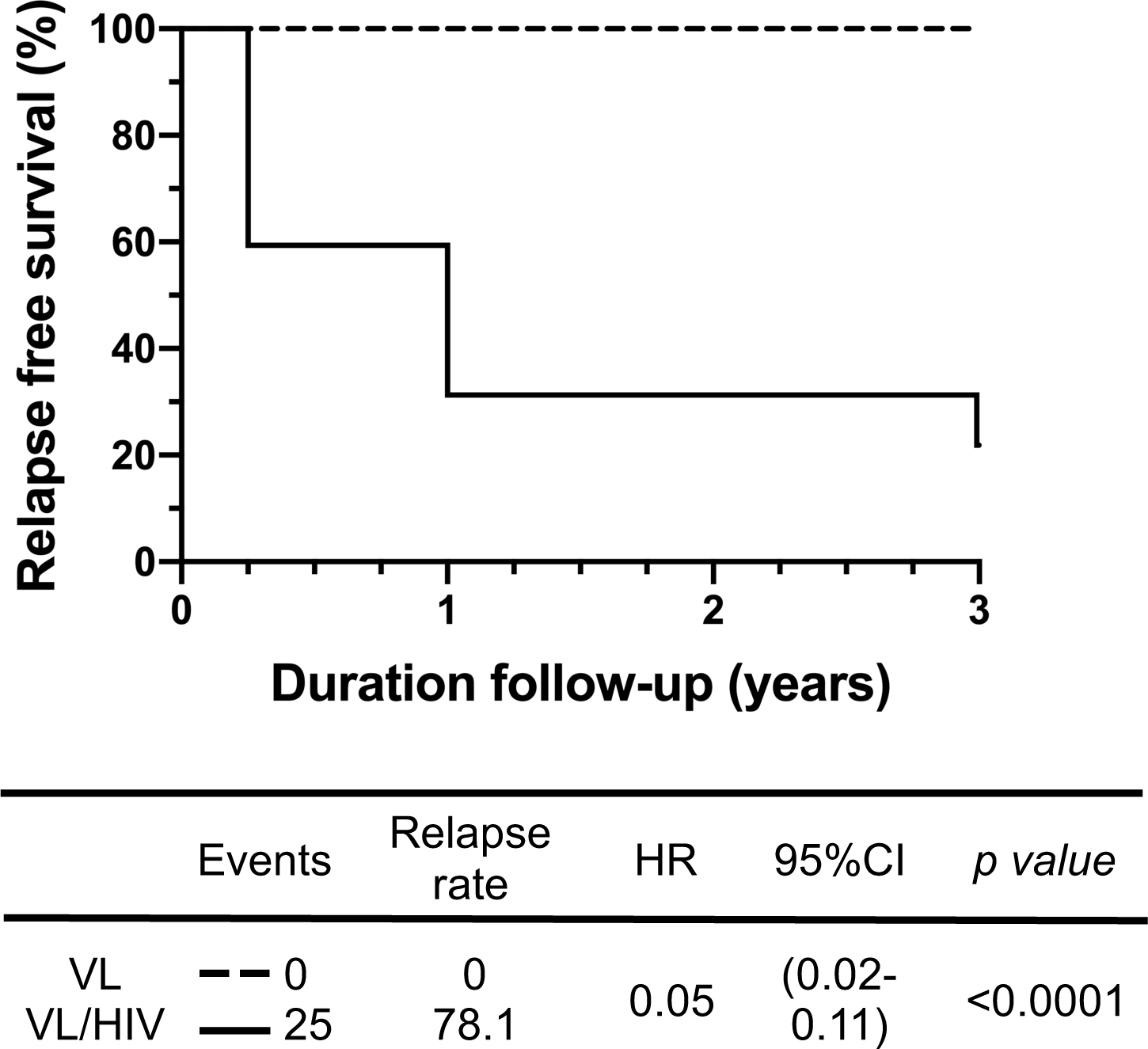
Relapse free survival: Kaplan-Meier curves of participant VL relapses comparing VL to VL/HIV patients. The hazard ratios (with 95% confidence intervals and *p* values) obtained from the Cox model indicated the change in survival following treatment for VL for these groups. HR, hazard ration; CI, confidence interval.

The 46 VL patients who reached the end of the anti-leishmanial treatment responded well to their first course of treatment. In the VL/HIV group, 25 patients responded well to the first course of anti-leishmanial treatment and had a negative test of cure at EoT and were therefore discharged from the hospital. 16 VL/HIV patients responded poorly to the first line of treatment and had a positive TOC at EoT, they therefore required a longer course of anti-leishmanial drugs. We compared the duration of this initial treatment between VL/HIV patients who relapsed and those who didn’t relapse during follow-up: results presented in Figure S1A show that the duration of treatment was similar in both groups of coinfected patients.

### Parasite grades and viral load

We first compared parasite grades as measured in splenic aspirates in VL and VL/HIV patients (24). As shown in Figure 2A, parasite grades in VL/HIV patients at time of diagnosis (ToD) were significantly higher than those in VL patients (*p*<0.001); despite a similar duration of symptoms by patients in both groups (VL: 2.0±0.2 vs VL/HIV: 2.0±0.2 months, *p*>0.05, Figure S2). The median parasite grade in VL/HIV patients at time of relapse was similar to that of VL/HIV patients at ToD (6.0±0.4, p>0.05, data not shown). Since parasite grade is mainly measured at ToD, when the spleen is easily palpable, we used RNAseq to measure the total expression of *L. donovani* mRNAs (*Ld* mRNA) in blood (Figure 2B). In agreement with the parasite grades measured at ToD, there was significantly more *Ld* mRNA in VL/HIV patients; these levels decreased significantly in both cohorts of patients after anti-leishmanial treatment at EoT but stayed significantly higher in VL/HIV patients. VL/HIV patients who relapsed during follow-up displayed significantly higher total expression of *Ld* mRNAs in blood than to those without relapse (Figure 2C).

**Figure 2:**
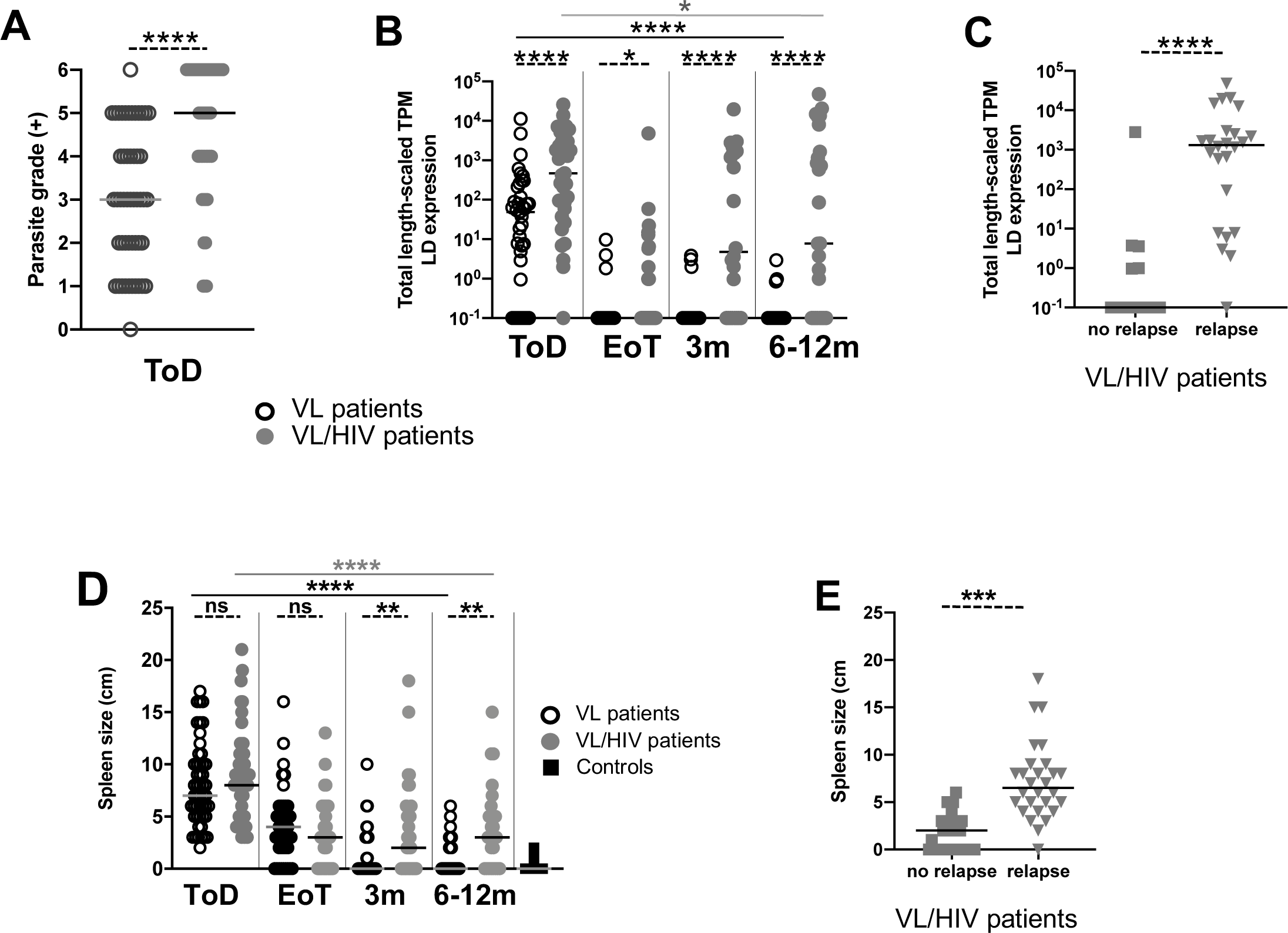
Parasite load and spleen size: **A.** Quantification of *Leishmania* amastigotes in smears of splenic aspirates collected from VL (n=50) and VL/HIV (n=49) patients at ToD. **B.** Quantification of the total expression of *L. donovani* mRNA in blood from VL (ToD: n=40, EoT: n=31, 3m: n=31, 6-12m: n=37) and VL/HIV (ToD: n=35, EoT: n=30, 3m: n=24, 6-12m: n=25) patients. **C.** Quantification of the total *L. donovani* mRNA expression in blood from VL/HIV who did not relapse (n=13) and who relapse (n=24) after successful anti-leishmanial treatment. **D** Spleen size as measured in cm below the costal margin on VL (ToD: n=50, EoT: n=45, 3m: n=36, 6-12m: n=26), VL/HIV (ToD: n=49, EoT: n=39, 3m: n=32, 6-12m: n=26) patients and controls (n=25). **E.** Spleen size as measured in cm below the costal margin on VL/HIV patients who did not relapse (n=28) and those who relapsed (n=37) after successful anti-leishmanial treatment. Each symbol represents the value for one individual, the straight lines represent the median. Statistical differences between VL and VL/HIV patients at each time point or between no relapse and relapse were determined using a Mann-Whitney test; statistical differences between the 4 different time points for each cohort of patients were determined by Kruskal-Wallis test. LD mRNA= *L. donovani* mRNA. ToD=Time of Diagnosis; EoT=End of Treatment; 3m=3 months post EoT; 6-12m=6-12 months post EoT. ns=not significant.

Measurement of plasma HIV-1 viral load showed that despite being on ART, 58.9% of VL/HIV patients still had detectable viral loads (Table 1). There were no significant differences in either viral load (Table 1A) or total expression of HIV mRNA between time points (Table 1B) or between patients with and without relapse during follow-up (Table 1C). There was no correlation between *L. donovani* mRNAs and viral loads in VL/HIV patients who relapsed during follow-up (*p*=0.3356, data not shown); similar results were obtained with the correlation between the total expression of HIV-1 and *L. donovani* mRNAs (*p*=0.0745, data not shown). Of note, there was no systematic pattern in how viral loads varied through follow-up (Figure S3).

**Table 1:** Viral load. **A.** HIV-1 viral load in plasma from VL/HIV (ToD: n=39, EoT: n=33, 3m: n=27, 6-12m: n=21) patients. **B.** Quantification of the total expression of HIV mRNA in blood from VL/HIV patients (ToD: n=35, EoT: n=30, 3m: n=24, 6-12m: n=25). **C.** Quantification of the total HIV mRNA expression in blood from VL/HIV who did not relapse (n=13) and who relapsed (n=24) after successful anti-leishmanial treatment (3 and 6-12 months). Statistical difference between the 4 different time points was determined by Kruskal-Wallis test and between relapse and no relapse by Mann-Whitney test. ToD=Time of Diagnosis; EoT=End of Treatment; 3m=3 months post EoT; 6-12m=6-12 months post EoT.

|  | **ToD** | **EoT** | **3m** | **6-12m** | **p value** |
| --- | --- | --- | --- | --- | --- |
| **VL/HIV** | 255±402369 | 150±115.951 | 0.1±61.395 | 0.1±200767 | 0.3632 |

**B. Viral load HIV mRNA**
| **HIV mRNA** | **ToD** | **EoT** | **3m** | **6-12m** | **p value** |
| --- | --- | --- | --- | --- | --- |
| **VL/HIV** | 673.0±492848 | 150.0±62366 | 75.±81420 | 273.5±141025 | 0.4205 |

**C. Viral load HIV mRNA in VL/HIV patients with and without relapse**
| **HIV mRNA** | **no relapse** | **relapse** | **p value** |
| --- | --- | --- | --- |
| **VL/HIV** | 0.1±154.1 | 397.1±97202 | 0.0575 |

### Clinical presentation

At ToD, a range of clinical and laboratory data are collected from each patient before the start of anti-leishmanial therapy.

#### Fever

results presented in Table 2 show that whereas both VL and VL/HIV patients had increased body temperature at ToD (controls: 36.0±0.1°C, *p*<0.0001), it was significantly lower in VL/HIV patients (*p*=0.023); and decreased over time in both groups (*p*<0.0001) but at 6-12 months (6-12m) was higher in VL/HIV patients as compared to VL patients (*p*=0.023). There was no significant difference in the body temperature of VL/HIV patients with and without relapse during follow-up (data not shown).

**Table 2:**
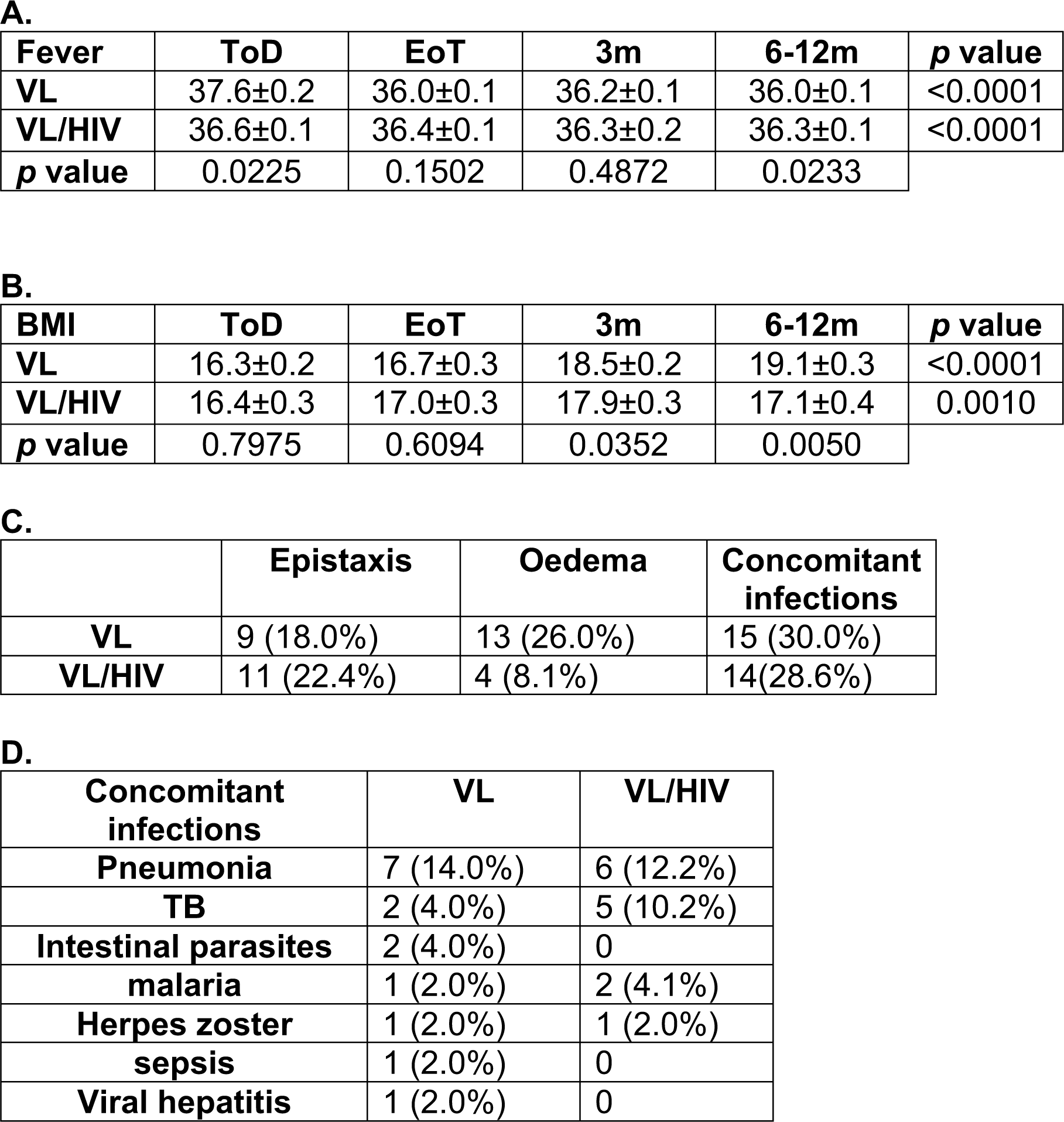
Clinical symptoms A. **A.** Body temperature was measured on VL (ToD: n=49, EoT: n=45, 3m: n=36, 6-12m: n=26), VL/HIV (ToD: n=49, EoT: n=38, 3m: n=32, 6-12m: n=26) patients and controls (n=25). **B.** BMI of VL (ToD: n=49, EoT: n=44, 3m: n=36, 6-12m: n=25), VL/HIV (ToD: n=49, EoT: n=39, 3m: n=32, 6-12m: n=26) patients and controls (n=25). **C**. Numbers and percentages of VL and VL/HIV patients presenting at ToD with Epistaxis, oedema or concomitant infections. **D.** Numbers and percentages of VL and VL/HIV patients presenting with the different concomitant infections. Statistical differences between VL and VL/HIV patients at each time point were determined using a Mann-Whitney test and statistical differences between the 4 different time points for each cohort of patients were determined by Kruskal-Wallis test. ToD=Time of Diagnosis; EoT=End of Treatment; 3m=3 months post EoT; 6-12m=6-12 months post EoT. ns=not significant.

| **Fever** | **ToD** | **EoT** | **3m** | **6-12m** | **p value** |
| --- | --- | --- | --- | --- | --- |
| **VL** | 37.6±0.2 | 36.0±0.1 | 36.2±0.1 | 36.0±0.1 | <0.0001 |
| **VL/HIV** | 36.6±0.1 | 36.4±0.1 | 36.3±0.2 | 36.3±0.1 | <0.0001 |
| **p value** | 0.0225 | 0.1502 | 0.4872 | 0.0233 |  |

**B.**
| **BMI** | **ToD** | **EoT** | **3m** | **6-12m** | **p value** |
| --- | --- | --- | --- | --- | --- |
| **VL** | 16.3±0.2 | 16.7±0.3 | 18.5±0.2 | 19.1±0.3 | <0.0001 |
| **VL/HIV** | 16.4±0.3 | 17.0±0.3 | 17.9±0.3 | 17.1±0.4 | 0.0010 |
| **p value** | 0.7975 | 0.6094 | 0.0352 | 0.0050 |  |

**C.**
|  | **Epistaxis** | **Oedema** | **Concomitant infections** |
| --- | --- | --- | --- |
| **VL** | 9 (18.0%) | 13 (26.0%) | 15 (30.0%) |
| **VL/HIV** | 11 (22.4%) | 4 (8.1%) | 14 (28.6%) |

**D.**
| **Concomitant infections** | **VL** | **VL/HIV** |
| --- | --- | --- |
| **Pneumonia** | 7 (14.0%) | 6 (12.2%) |
| **TB** | 2 (4.0%) | 5 (10.2%) |
| **Intestinal parasites** | 2 (4.0%) | 0 |
| **malaria** | 1 (2.0%) | 2 (4.1%) |
| **Herpes zoster** | 1 (2.0%) | 1 (2.0%) |
| **sepsis** | 1 (2.0%) | 0 |
| **Viral hepatitis** | 1 (2.0%) | 0 |

#### Hepatosplenomegaly

As shown in Figure 2D spleen sizes were similarly increased in both cohorts of patients at ToD and decreased at EoT. These continued to decrease in VL patients but stayed higher in VL/HIV patients at 3 months (3m) and 6-12m. Furthermore, during follow-up of VL/HIV patients, spleens were significantly more enlarged in VL/HIV patients who relapsed (Figure 2E). The liver was also palpable at ToD in 40.0% of VL and 63.3% of VL/HIV patients and was significantly more enlarged in VL/HIV as compared to VL patients (0.0±0.5 and 3.0±0.5 cm, respectively, *p*=0.012, Figure S4); the liver size decreased significantly at EoT (Figure S4). There was no significant difference in liver size in VL/HIV patients with and without relapse during follow-up (data not shown).

#### Body mass index (BMI)

The median BMI of patients with VL and VL/HIV was below at ToD (VL patients: 16.4±0.3, VL/HIV patients: 16.7±0.3, Table 2B). The BMI of VL patients increased over time and at 3m, was similar to those of controls (controls: 19.9±0.6, p>0.05); in contrast, the BMI of the VL/HIV patients stayed significantly lower compared to VL patients and controls over time (Table 2B).

Epistaxis, oedema and concomitant infections are clinical symptoms that are routinely recorded at ToD in these patients. As shown in Table 2C, there were no significant differences between the number of patients experiencing epistaxis or co-infections (pneumonia, TB, intestinal parasites, malaria, herpes zoster, sepsis or viral hepatitis, Table 2D), but there were more VL patients presenting with oedema.

Liver and kidney function are also routinely measured at ToD and EoT. Results presented in Table 3 show significantly lower levels of SGOT and SGPT in the VL/HIV groups at both ToD and EoT. The levels of BUN and creatinine decreased significantly in the VL group at EoT but remained similar in the VL/HIV groups.

**Table 3:** Liver and kidney function tests. Liver and kidney functions were measured in the plasma of VL patients as described in Materials and Methods. SGPT=serum glutamic oxaloacetic transaminase, SGOT= serum glutamic pyruvic transaminase, and BUN=blood urea nitrogen. Statistical difference was determined by a Mann-Whitney test The values in italic and in parenthesis represent the normal values. ToD=Time of Diagnosis; EoT=End of Treatment

| **SGPT**<br>**(*<43 U/L*)** | **ToD** | **EoT** | ***p* value** |
| --- | --- | --- | --- |
| **VL** | 34.0 ± 7.5 | 43.5 ± 5.7 | ns |
| **VL/HIV** | 20.0 ± 2.1 | 28.5 ± 4.2 | ns |
| ***p* value** | <0.001 | 0.002 |  |

| **SGOT**<br>**(*<38 U/L*)** | **ToD** | **EoT** | ***p* value** |
| --- | --- | --- | --- |
| **VL** | 62.0 ± 13.7 | 61.0 ± 5.2 | ns |
| **VL/HIV** | 41.0 ± 4.9 | 35.0 ± 5.6 | ns |
| ***p* value** | <0.001 | <0.001 |  |

| **BUN**<br>**(*4.7-23.5 mg/dL*)** | **ToD** | **EoT** | ***p* value** |
| --- | --- | --- | --- |
| **VL** | 12.0 ± 1.9 | 9.0 ± 0.5 | 0.0022 |
| **VL/HIV** | 12.4 ± 1.1 | 13.8 ± 1.1 | ns |
| ***p* value** | ns | <0.001 |  |

| **Creatinine**<br>**(*0.6-1.1mg/dL*)** | **ToD** | **EoT** | ***p* value** |
| --- | --- | --- | --- |
| **VL** | 0.9 ± 0.1 | 0.8 ± 0.1 | 0.0092 |
| **VL/HIV** | 0.9 ± 0.2 | 1.0 ± 0.1 | ns |
| ***p* value** | ns | <0.001 |  |

### Haematological profile

White and red blood cell and platelet counts were all significantly decreased at ToD (Figure 3A, C and E), except for platelet counts that were higher in VL/HIV patients (Figure 3E). White blood cell counts increased in both groups at EoT, but thereafter only continued to increase in VL, but not in VL/HIV patients. Of note, both VL and VL/HIV patients had not restored levels of white blood cells 6-12m post EoT. Similarly, red cell and platelet counts increased in VL patients, and, in contrast to the counts in VL/HIV patients, were restored to levels similar to controls at EoT. Furthermore, during follow-up of VL/HIV patients, white and red blood cell and platelet counts were significantly lower in VL/HIV patients who relapsed.

**Figure 3:**
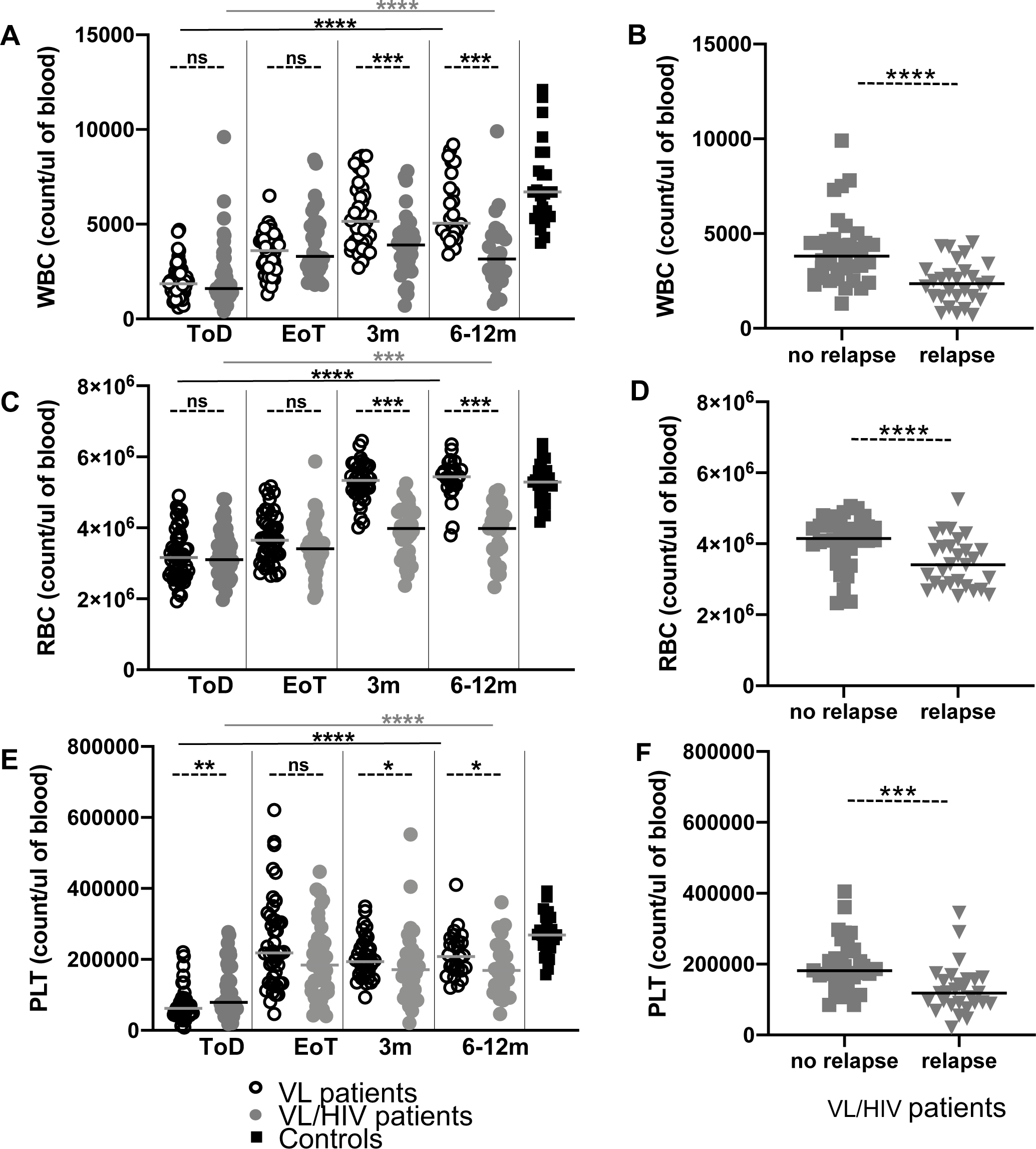
Haematological data: **A.** White blood cells, **C.** Red blood cells and **E.** Platelet counts in the blood of VL (ToD: n=50, EoT: n=45, 3m: n=36, 6-12m: n=26), VL/HIV (ToD: n=49, EoT: n=39, 3m: n=32, 6-12m: n=27) patients and controls (n=25). **B.** White blood cells, **D.** Red blood cells and **F.** Platelet counts in the blood of VL/HIV patients who did not relapse (n=34) and those who relapsed (n=28) after successful anti-leishmanial treatment. Each symbol represents the value for one individual, the straight lines represent the median. Statistical differences between VL and VL/HIV patients at each time point or between no relapse and relapse were determined using a Mann-Whitney test and statistical differences between the 4 different time points for each cohort of patients were determined by Kruskal-Wallis test. ToD=Time of Diagnosis; EoT=End of Treatment; 3m=3 months post EoT; 6-12m=6-12 months post EoT. ns=not significant.

In summary, the results from the clinical data show that over time, VL/HIV patients maintained high parasite loads, hepatosplenomegaly and low BMI, and remained pancytopenic.

## IMMUNOLOGICAL DATA

### Antigen-specific production of IFNγ and IL-10 by whole blood cells from VL and VL/HIV patients

Using a whole blood assay (WBA), we have previously shown that whole blood cells (WBC) from VL patients from Northern Ethiopia displayed an impaired capacity to produce IFNγ in response to stimulation with soluble *Leishmania* antigens (SLA) at ToD; but that these cells gradually regained their capacity to produce IFNγ over time after successful treatment (18). Results presented in Figure 4A show that antigen-specific production of IFNγ was low at ToD in both cohorts of patients, but increased significantly in VL patients at EoT, and was restored during follow-up. In contrast, the levels of IFNγ produced by WBC from VL/HIV patients remained low at all time points. This was also true in longitudinal follow-up of patients (figures S5A and B, *p*=0.0038 and *p*=0.1682, respectively). We also compared the levels of antigen-specific IFNγ in VL/HIV patients with and without relapse: in the longitudinal follow-up (Figure S5), those who did not relapse produced significantly more IFNγ (p=0.0022); when assessed cross-sectionally (Figure 4B), the median levels of IFNγ produced by WBC from patients who did not relapse after treatment were also significantly higher as compared to those who relapsed and were similar to those measured at ToD in VL patients (Figure 4A).

**Figure 4:**
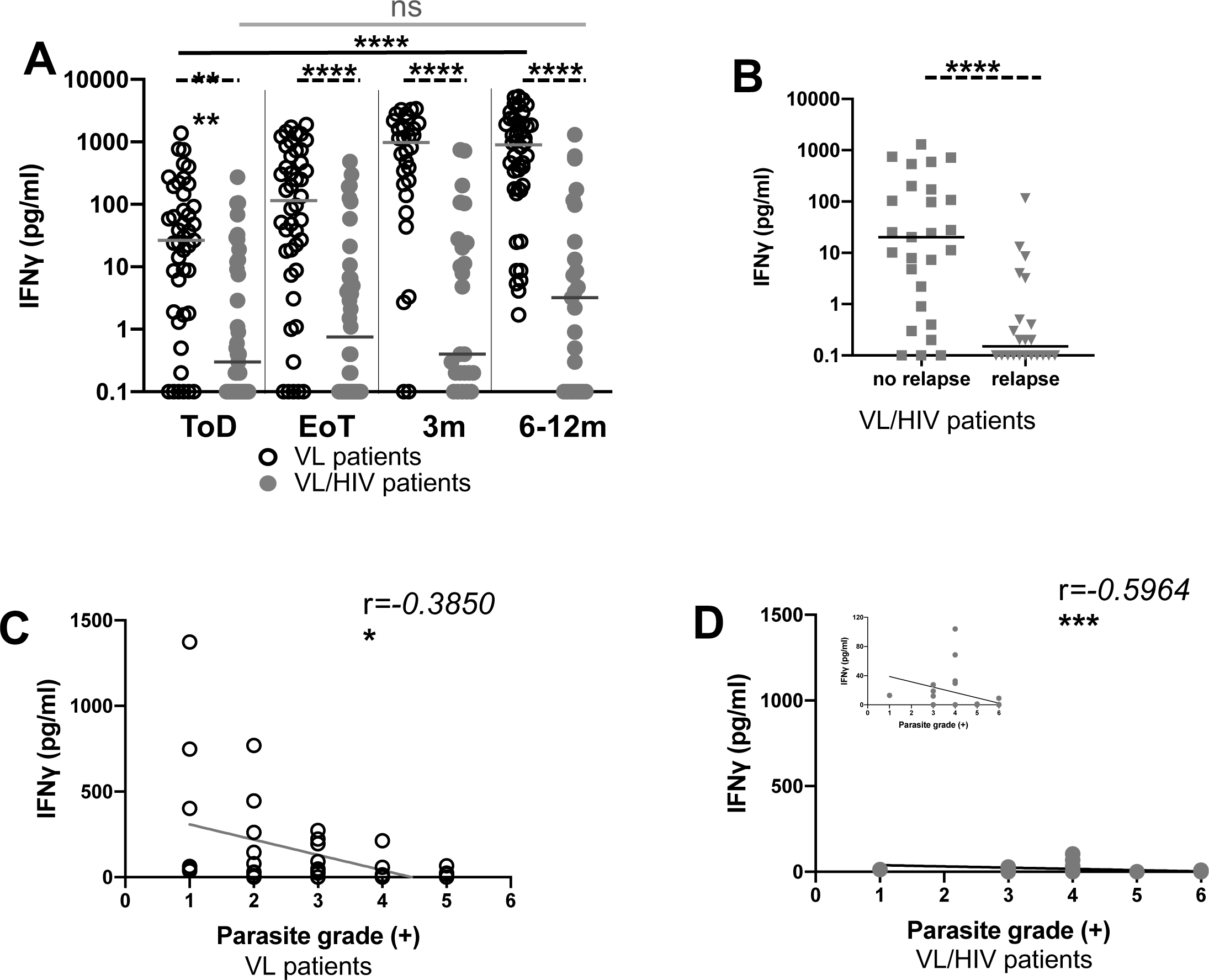
Whole blood assay: antigen-specific production of IFNγ: **A.** Whole blood cells from VL (ToD: n=43, EoT: n=44, 3m: n=30, 6-12m: n=44) and VL/HIV patients (ToD: n=39, EoT: n=40, 3m: n=25, 6-12m: n=25) were cultured in the presence of SLA and IFN*γ* was measured by ELISA in the supernatant after 24hrs. **B.** Comparison of the levels of antigen-specific IFN*γ* produced by whole blood cells from VL/HIV patients who did not relapse (n=27) and those who relapsed (n=22) after successful anti-leishmanial treatment. **C.** Correlation between parasite grades and IFNγ at ToD in VL patients (n=38) and **D.** VL/HIV patients (n=34). Each symbol represents the value for one individual, the straight lines represent the median. Statistical differences between VL and VL/HIV patients at each time point or between no relapse and relapse were determined using a Mann-Whitney test; statistical differences between the 4 different time points for each cohort of patients were determined by Kruskal-Wallis test and the correlation by Spearman’s rank test. ToD=Time of Diagnosis; EoT=End of Treatment; 3m=3 months post EoT; 6-12m=6-12 months post EoT. ns=not significant.

There was a clear correlation between IFNγ concentrations and parasite grade at ToD (Figures 4C and D).

IFNγ production in response to phytohemagglutinin (PHA) followed the same pattern as antigen-specific stimulation, failing to recover from a low level in the VL/HIV patients (Tables 4).

**Table 4A:** Production of IFNɣ in response to SLA and PHA.

| **VL** | **ToD** | **EoT** | **3m** | **6-12m** | **p value** |
| --- | --- | --- | --- | --- | --- |
| **SLA** | 28.4±41.3 | 115.4±80.4 | 972.2±205.0 | 898.4±229.6 | <0.0001 |
| **PHA** | 30.6±35.6 | 177.8±123.0 | 190.8±186.0 | 389.5±196.7 | 0.0001 |
| **p value** | 0.6728 | 0.6552 | 0.0637 | 0.1971 |  |

| **VL/HIV** | **ToD** | **EoT** | **3m** | **6-12m** | **p value** |
| --- | --- | --- | --- | --- | --- |
| **SLA** | 0.3±7.8 | 0.8±15.5 | 0.4±40.5 | 3.2±58.4 | 0.3411 |
| **PHA** | 6.4±65.9 | 33.1±129.1 | 14.5±101.1 | 14.2±148.6 | 0.5850 |
| **p value** | 0.0001 | 0.0002 | 0.0249 | 0.0524 |  |

**Table 4B:** Production of IFNɣ in response to PHA (VL and VL/HIV patients) A. Whole blood cells from VL (ToD: n=43, EoT: n=44, 3m: n=30, 6-12m: n=44) and VL/HIV patients (ToD: n=39, EoT: n=40, 3m: n=25, 6-12m: n=25) were cultured in the presence of SLA and PHA and IFN*γ* levels in the supernatants were measured by ELISA after 24hrs. Statistical differences between VL and VL/HIV patients at each time point were determined using a Mann-Whitney test; statistical differences between the 4 different time points for each cohort of patients were determined by Kruskal-Wallis test B. Comparison of the levels of IFNɣ produced in response to PHA between VL and VL/HIV patients. Statistical were determined using a Mann-Whitney test ToD=Time of Diagnosis; EoT=End of Treatment; 3m=3 months post EoT; 6-12m=6-12 months post EoT.

| **PHA** | **ToD** | **EoT** | **3m** | **6-12m** |
| --- | --- | --- | --- | --- |
| **VL** | 30.6±35.6 | 177.8±123.0 | 190.8±186.0 | 389.5±196.7 |
| **VL/HIV** | 6.4±65.9 | 33.1±129.1 | 14.5±101.1 | 14.2±148.6 |
| **p value** | 0.1089 | 0.0764 | 0.0075 | <0.0001 |

We have previously shown that in VL patients in Ethiopia, antigen-specific production of IL-10 was low in the WBA (18). Results presented in Table 5A confirm our previous data and show that the levels of IL-10 remained low over time (Table 5A).

**Table 5A:** Production of IL-10 in response to SLA and PHA.

| **VL** | **ToD** | **EoT** | **3m** | **6-12m** | **p value** |
| --- | --- | --- | --- | --- | --- |
| **SLA** | 2.9±1.3 | 0.1±0.76 | 1.7±1.3 | 0.3±3.1 | 0.0153 |
| **PHA** | 26.9±19.4 | 142.1±30.2 | 648.9±67.4 | 390.7±38.7 | <0.0001 |
| **p value** | 0.0002 | <0.0001 | <0.0001 | <0.0001 |  |

| **VL/HIV** | **ToD** | **EoT** | **3m** | **6-12m** | **p value** |
| --- | --- | --- | --- | --- | --- |
| **SLA** | 0.3±3.1 | 2.6±1.8 | 0.7±0.7 | 0.2±1.2 | 0.1524 |
| **PHA** | 25.4±16.9 | 227.3±42.0 | 319.2±41.6 | 255.5±56.3 | <0.0001 |
| **p value** | <0.0001 | <0.0001 | <0.0001 | <0.0001 |  |

In contrast, the production of IL-10 in response to PHA increased overtime in both VL and VL/HIV patients (Table 5A) and was systematically higher as compared to SLA in both cohorts (Table 5A). The levels of IL-10 produced in response to PHA were similar in VL and VL/HIV at ToD, EoT and 6-12m (Table 5B). Sustained levels of antigen-specific IL-10 production by WBC are thus not associated with disease severity.

**Table 5B:**
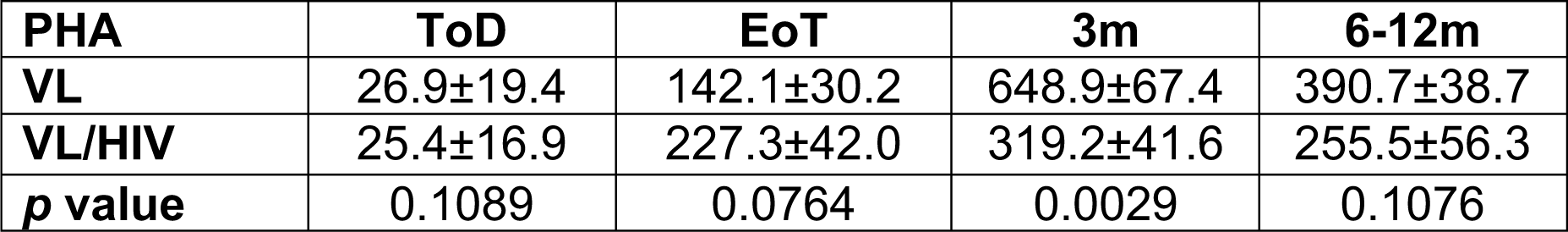
Production of IL-10 in response to PHA (VL and VL/HIV patients) **A.** Whole blood cells from VL (ToD: n=43, EoT: n=44, 3m: n=30, 6-12m: n=44) and VL/HIV patients (ToD: n=39, EoT: n=40, 3m: n=25, 6-12m: n=25) were cultured in the presence of SLA and PHA and IL-10 levels ELISA in the supernatant were measured by after 24hrs. Statistical differences between VL and VL/HIV patients at each time point were determined using a Mann-Whitney test; statistical differences between the 4 different time points for each cohort of patients were determined by Kruskal-Wallis test **B.** Comparison of the levels of IL-10 produced in response to PHA between VL and VL/HIV patients. Statistical were determined using a Mann-Whitney test ToD=Time of Diagnosis; EoT=End of Treatment; 3m=3 months post EoT; 6-12m=6-12 months post EoT.

### CD4^+^ T cell counts in VL and VL/HIV co-infected patients

To determine whether the low production of IFNγ is associated with low T cell counts, the absolute CD4^+^ T cell counts were measured in both cohorts of patients. CD4^+^ T cell counts in VL patients were low at ToD but were restored at EoT (controls CD4^+^ T cell counts: 455±16.4 cells/μl of blood, data not shown). Despite an increase in CD4^+^ T cell counts at EoT in VL/HIV co-infected patients, CD4^+^ T cell counts remained significantly lower at all time points (Figure 5A). Longitudinal follow-up of individual patients also shows a significant increase over time in CD4^+^ T cell counts in VL, but not in VL/HIV patients (Figure S6 A and B, *p*=0.0069 and *p*=0.0923, respectively). Median CD4^+^ T cell counts in patients who did not relapse after treatment was significantly higher as compared to those who relapsed (Figure 5B).

**Figure 5:**
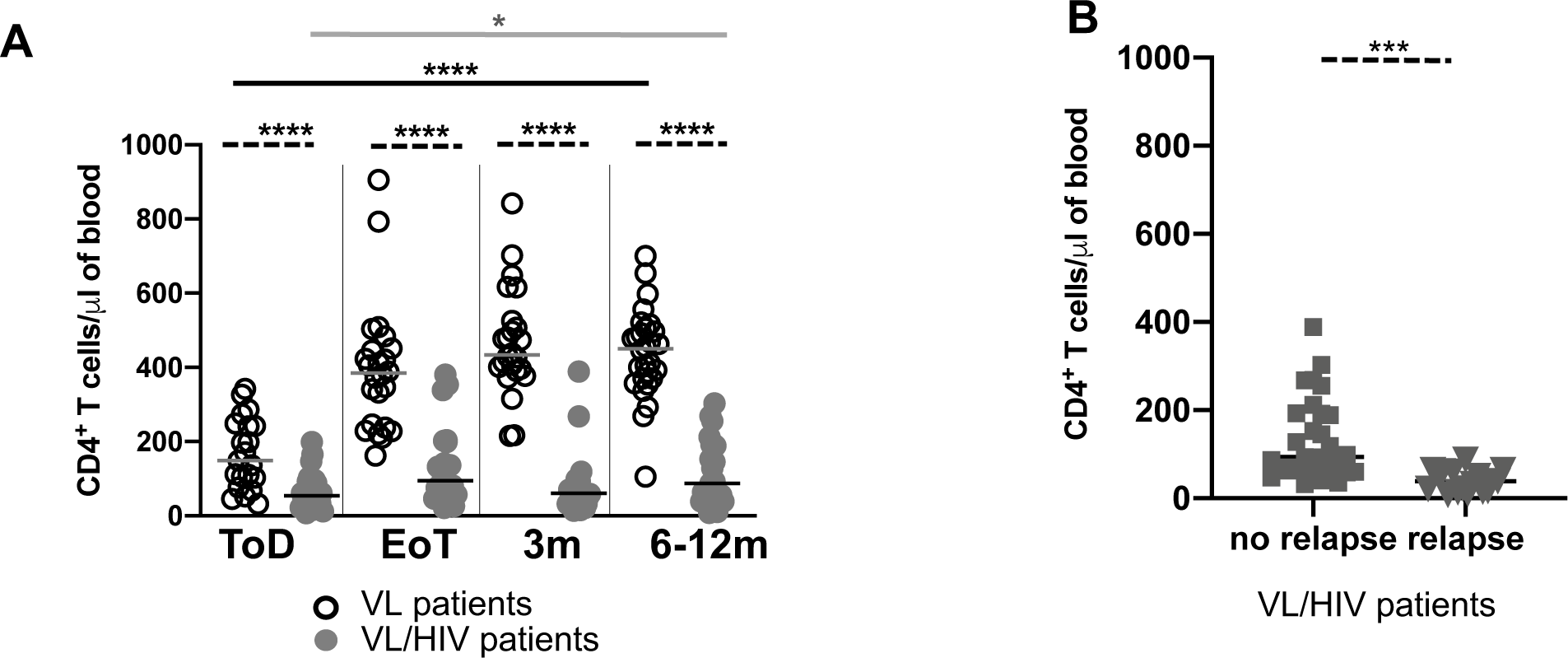
CD4^+^ T cell counts: **A.** CD4^+^ T cell counts were measured by flow cytometry in the blood of VL (ToD: n=21, EoT: n=24, 3m: n=24, 6-12m: n=28) and VL/HIV patients (ToD: n=27, EoT: n=24, 3m: n=21, 6-12m: n=25). **B.** Comparison of CD4^+^ T cell counts in VL/HIV patients who did not relapse (n=29) and those who relapsed (n=17). Each symbol represents the value for one individual, the straight lines represent the median. Statistical differences between VL and VL/HIV patients at each time point or between no relapse and relapse were determined using a Mann-Whitney test and statistical differences between the 4 different time points for each cohort of patients were determined by Kruskal-Wallis test. ToD=Time of Diagnosis; EoT=End of Treatment; 3m=3 months post EoT; 6-12m=6-12 months post EoT. ns=not significant.

### Inflammatory cytokines in VL and VL/HIV patients

It is well established that markers of inflammation are high in the plasma of VL patients (19), and indeed, the levels of CRP were elevated at ToD and fell to physiological levels at the end of the follow-up (Figure S7)(25). In contrast, in VL/HIV patients, these levels decreased over time, but remained significantly higher (Figure S7). Furthermore, results presented in Figures 6A, B and C show that the levels of TNF*α*, IL-8 and IL-6 were high in the plasma of both cohorts of patients at ToD. These levels decreased in both groups at EoT, and continue to decrease over time in the VL, but not in the VL/HIV cohort. The levels IFNγ were significantly higher at ToD in the plasma of VL patients and dropped to levels similar to those of VL/HIV patients during the follow-up (Figure 6D). IL-1*β* was only detectable at ToD in the VL group and dropped to levels below detection limit at EoT and IL-12 levels were below detection limits in the large majority of the plasma (data not shown). IL-10 levels were high at ToD in the plasma of VL patients and decreased at EoT, and at both time points, remained significantly higher as compared to the VL/HIV group (Figure 6E); and continued to decrease in the VL group after 3 and 6-12 months but remained significantly higher in the VL/HIV group (Figure 6E). The levels of the anti-inflammatory cytokines IL-4 and IL-13 were largely below detection limits (data not shown).

**Figure 6:**
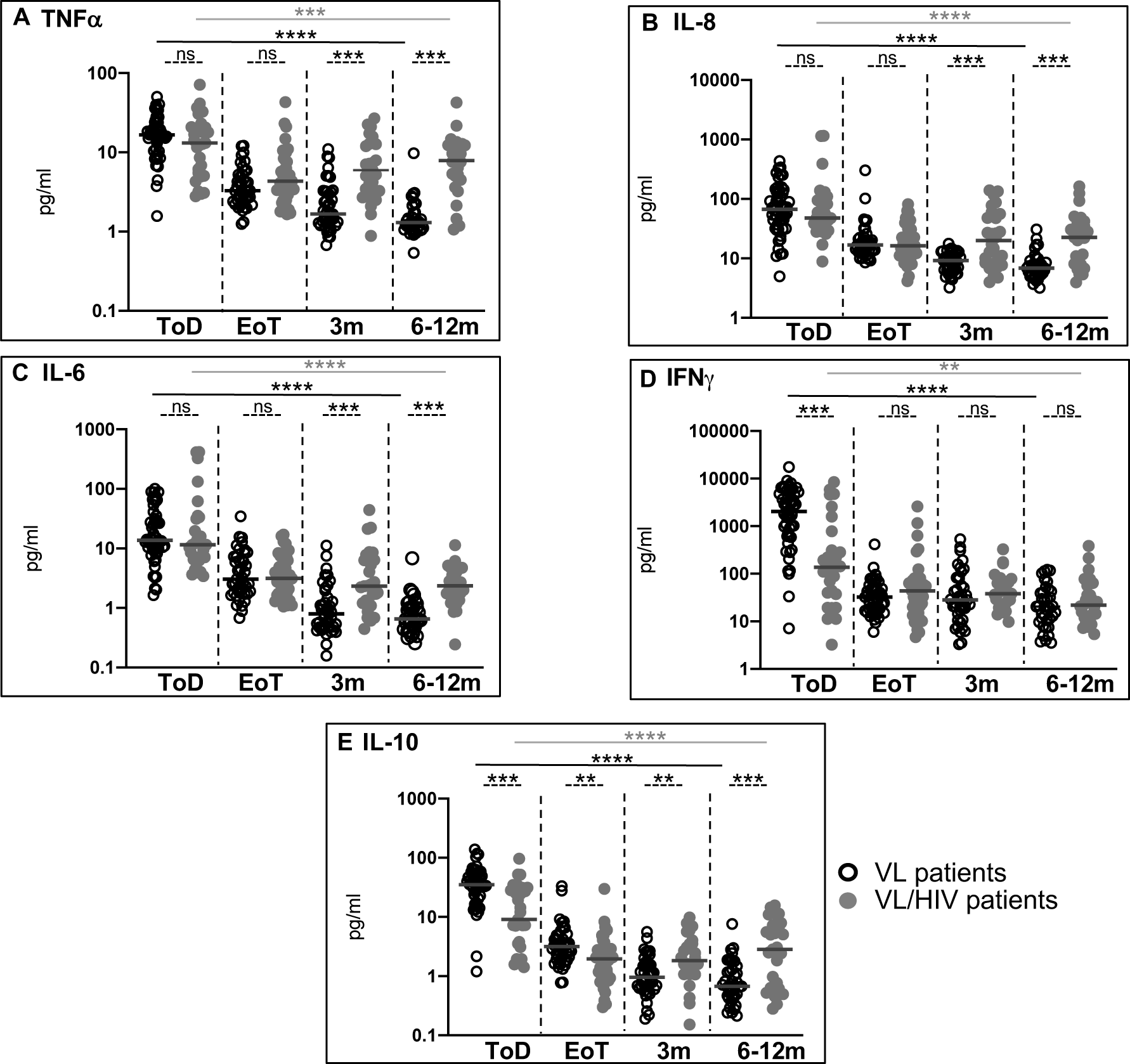
Cytokines in the plasma of VL and VL/HIV patients: Levels of **A.** TNF*α*; **B.** IL-8, **C.** IL-6; **D.** IFN*γ* and **E.** IL-10 were measured in the plasma isolated from the blood of VL (ToD: n=42, EoT: n=33, 3m: n=32, 6-12m: n=27), VL/HIV patients (ToD: n=29, EoT: n=39, 3m: n=28, 6-12m: n=28) and controls (n=22) by multiplex assay. Each symbol represents the value for one individual, the straight lines represent the median. Statistical differences between VL and VL/HIV patients at each time point or between no relapse and relapse were determined using a Mann-Whitney test and statistical differences between the 4 different time points for each cohort of patients were determined by Kruskal-Wallis test. ToD=Time of Diagnosis; EoT=End of Treatment; 3m=3 months post EoT; 6-12m=6-12 months post EoT. ns=not significant.

### Inhibitory receptors expressed by CD4^+^ T cells from VL and VL/HIV co-infected patients

Immune checkpoint molecules can trigger immunosuppressive signalling pathways that can drive T cells into a state of hyporesponsiveness, with reduced effector functions and sustained expression levels of inhibitory receptors (26). Here we focused on CTLA-4 and PD1 expression on CD4^+^ T cells. CTLA-4 expression on T cells from VL, VL/HIV patients and controls was below detection limit (<1%, data not shown). Results presented in Figure 7A show that the PD1 iMFI of CD4^+^ T cells were high at ToD in both cohorts of patients, but decreased significantly in VL patients at the EoT, and were restored 3m after EoT (controls: CD4 PD1 iMFI: 12737±3095, p>0.05, data not shown). In contrast, CD4 PD1 iMFI in VL/HIV co-infected patients remained high after treatment and during follow-up (Figures 7A). The cross-sectional pattern was replicated in longitudinal trends for individual patients (Figures S8, *p*=0.0002 and *p*=0.0057, respectively). Next, we compared the levels of PD1 MFI during follow-up (3 to 6-12 months after EoT) in VL/HIV patients who relapsed and those who did not relapse: results presented in Figure 7B show that the expression levels of CD4 PD1 iMFI were significantly lower in VL/HIV patients who did not relapse after treatment.

**Figure 7:**
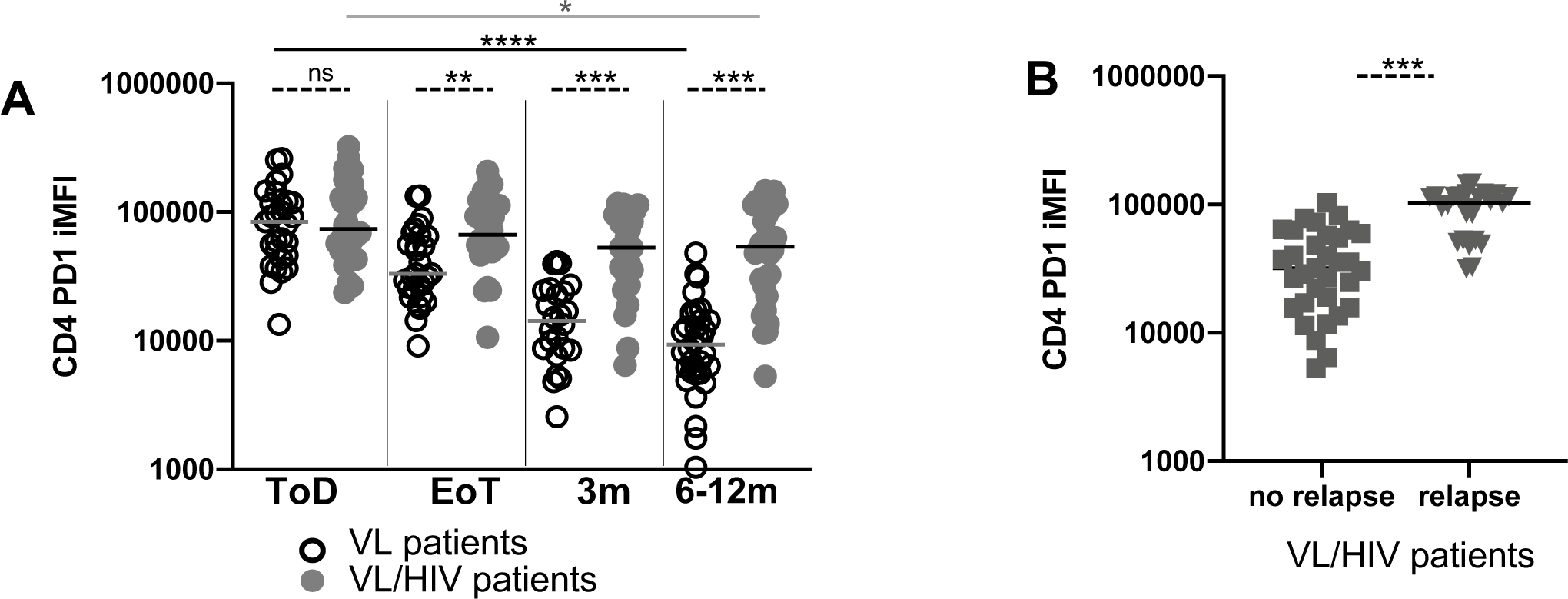
PD1 expression on CD4^+^ T cells. **A.** CD4 PD1 iMFI was measured by multiplying the % of CD4^+^ T cells and the median fluorescence intensity (MFI) of PD1 as measured by flow cytometry in the PBMCs of VL (ToD: n=29, EoT: n=29, 3m: n=24, 6-12m: n=34) and VL/HIV patients (ToD: n=28, EoT: n=32, 3m: n=26, 6-12m: n=32). **B.** Comparison of CD4 PD1 iMFI in the blood of VL/HIV patients who did not relapse (n=33) and those who relapsed (n=25). Each symbol represents the value for one individual, the straight lines represent the median. Statistical differences between VL and VL/HIV patients at each time point or between no relapse and relapse were determined using a Mann-Whitney test and statistical differences between the 4 different time points for each cohort of patients were determined by Kruskal-Wallis test. ToD=Time of Diagnosis; EoT=End of Treatment; 3m=3 months post EoT; 6-12m=6-12 months post EoT. ns=not significant.

These results show that cure and absence of subsequent relapse in VL patients is characterised by a significant decrease in the levels of PD1 expressed by CD4^+^ T cells at EoT. In contrast, CD4^+^ T cells from VL/HIV patients had persistently high expression of PD1. Furthermore, following successful treatment, CD4^+^ T cells from VL/HIV patients who did not relapse over time expressed significantly less PD1 than did those who relapsed.

In summary, our study has identified 3 immunological markers that are associated with VL relapse in VL/HIV patients: low CD4^+^ T cell count, low IFN*γ* production in the WBA, and high PD1 expression on CD4^+^ T cells.

### Prediction of relapse

Next, we formally tested whether the clinical and immunological parameters we measured at the end of treatment could be used to predict if and when the patients will relapse. Since none of the clinical parameters were significantly different at EoT between VL and VL/HIV patients, we tested whether the three immunological parameters we identified could be used as predictors of relapse. The results of the multinomial logistic regression analysis are summarised in Table 6.

**Table 6:** Multinomial regression to predict time of relapse in VL/HIV patients. Results of a multinomial logistic regression to predict relapse of VL/HIV individuals. CD4 (CD4+ T cell counts), IFN*γ* (WBA) and PD1 (CD4 PD1 iMFI) levels from EoT were used as predictors in predicting relapse at three potential time points; relapse at 3m, relapse at 6-12m, or no relapse recorded within 12m of EoT. *p<0.1; **p<0.05; ***p<0.01

|  | Dependent variable |  |
| --- | --- | --- |
|  | Relapse at 3m<br>(1) | Relapse at 6-12m<br>(2) |
| **CD4** | 0.071***<br>(0.000) | -0.164***<br>(0.000) |
| **IFNγ** | 0.044***<br>(0.000) | -1.321***<br>(0.000) |
| **PD1** | 0.0001***<br>(0.000) | 0.00003***<br>(0.00000) |
| **CD4:IFNγ** | 0.002***<br>(0.000) | 0.010***<br>(0.000) |
| **CD4:PD1** | 0.00000***<br>(0.00000) | 0.00000*<br>(0.00000) |
| **IFNγ:PD1** | 0.00000***<br>(0.00000) | 0.00000***<br>(0.00000) |
| **CD4:IFNγ:PD1** | 0.00000***<br>(0.000) | 0.00000***<br>(0.00000) |
| **Constant** | -9.553***<br>(0.000) | 8.897***<br>(0.000) |
| **Akaike Inf. Crit.** | 53.005 | 53.005 |

We trained a multinomial logistic regression model to predict early (at 3m follow-up), later (at 6-12m) or no relapse 12m follow-up, based on CD4 T cell counts, IFN*γ* and PD1 levels at EoT (full results in Table 6). This model performs well at distinguishing relapse times (Figure 8; Relapse at 3m AUC = 0.971, CI = 0.912 - 1.00; Relapse at 6-12m; AUC = 0.915, CI = 0.794 - 1.00) and particularly in distinguishing early relapse compared to relapse at later timepoints, which will be critical in identifying patients who need continuing treatment.

**Figure 8:**
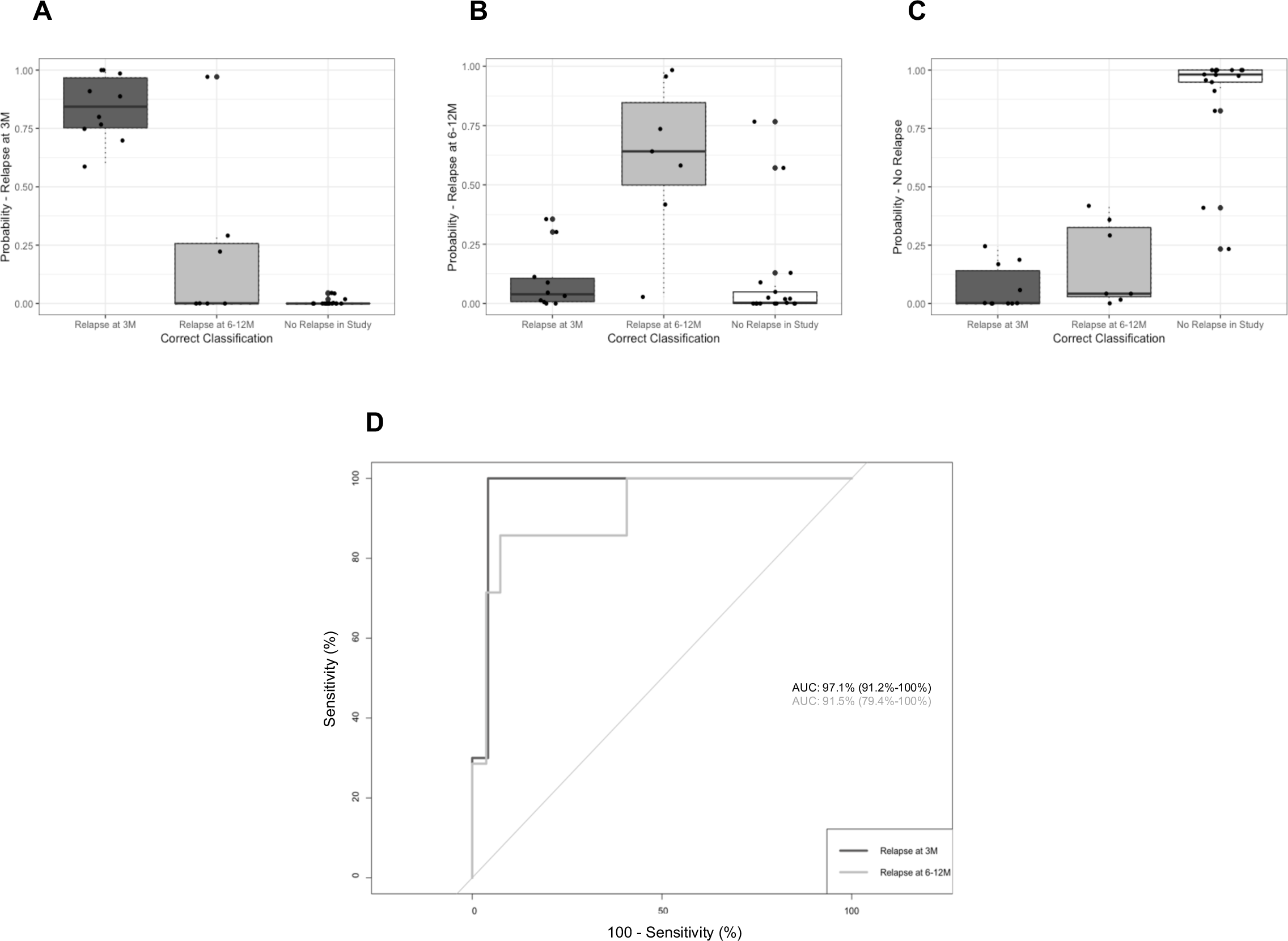
Performance of Multinomial Logistic Regression model in predicting relapse time for VL/HIV individuals. **A**. Performance of model in predicting VL/HIV individuals who do not relapse with VL within 12m of study EoT. **B.** Performance of model in predicting relapse at 6-12m from EoT. **C.** Performance of model in predicting VL relapse within 3m of EoT. **D**. ROC for predicting VL relapse at 3m and relapse at 6-12m groups for VL/HIV individuals from logistic regression model.

## DISCUSSION

The rate of VL/HIV coinfection is very high in Ethiopia and represents a major public health problem mainly due to the high rate of VL relapse in these patients. Currently there is no means to assess the probability of VL relapse after successful treatment. In this study we followed cohorts of VL and VL/HIV patients for up to 3 years, including detailed immunological follow-up for up to 12m. This detailed follow up reveals a higher rate of relapse than previously identified (12), with 78.1% of VL/HIV patients experiencing at least one relapse during the follow-up. Since nine VL/HIV patients were lost to follow-up in our study, it is possible that the relapse rate could be even higher. This higher relapse rate is likely to be due to the longer duration of our study and emphasizes the need for longer clinical follow-up of these patients.

Our results show that VL/HIV patients harbour a higher parasite load in splenic aspirates than VL patients at ToD. Quantifying parasite load in splenic aspirates over time in VL and VL/HIV patients has been an ongoing challenge, as it is not possible to perform splenic aspirates when the spleen size is <3cm below the costal margin. To evaluate the parasite loads in these patients over time, we measured the total expression of *Ld* mRNA in whole blood. This is the first study that compares parasites loads in VL and VL/HIV patients before and after treatment and over time. Parasite gene expression in blood was significantly correlated with spleen parasite load at ToD (p<0.0001, r=0.7597, data not shown) suggesting that measuring the total mRNA in whole blood is a good alternative to splenic aspirate. And indeed, a recent study showed a strong correlation between tissue parasite burden and blood parasite load measured by qPCR (27). While regular monitoring of parasite load is likely to be a valuable means to predict relapse, especially in VL/HIV patients, neither qPCR nor RNAseq are available in many of the leishmaniasis treatment centres in Ethiopia and therefore, alternative tests that can be performed in these settings are needed.

Despite being on ART, 58% of co-infected patients had detectable HIV viral load; 47 were on first line treatment and two on second line treatment. The high incidence of detectable viral load in this population could be due to: resistance to HIV drugs, which has been reported to both first and second line treatments (28); poor availability of ART for the population of migrant workers during the agricultural season; or due to poor adherence to antiretroviral therapy. While viral load in this population of HIV patients should be monitored more closely, VL/HIV patients who relapsed after successful anti-leishmanial treatment had similar viral load in plasma as those who did not relapse and there was no correlation between the viral load and total *Ld* mRNA expression. In agreement with previous studies, this suggests that VL relapse in VL/HIV patients occurs independently of the viral load (13, 29–31), but we cannot exclude that better long-term management of HIV in these patients might reduce the VL relapse rate.

As previously shown (32), both VL and VL/HIV patients have a low BMI at ToD. The follow-up of VL patients revealed that their BMI is only restored at the last time point, 6-12 months post the end of the anti-leishmanial treatment. In contrast, the BMI of VL/HIV patients remained below 18.5 throughout follow-up. It is generally accepted that malnutrition plays a key role in increased susceptibility to infection and/or disease severity by weakening both innate and acquired immunity (33, 34). It is therefore possible that a low BMI contributes to poor prognosis of VL/HIV patients.

The high levels of markers of inflammation (Figure 7) in their plasma might also contribute to their inability to regain weight as these cytokines have been shown to suppress appetite (35). Better management of malnutrition in these patients might also help to improve their ability to mount appropriate immune responses.

Severe pancytopenia is a hallmark of visceral leishmaniasis, it is thought to reflect bone marrow suppression and splenic sequestration. VL patients suffer from ineffective haematopoiesis; reticulo-endothelial cells infiltrate the bone marrow that becomes hyperplasic (36, 37) and *L. donovani* can also establish infection in the bone marrow. In addition to bone marrow failure, spleen sequestration is thought to play a major role in the severe pancytopenia observed in these patients (37, 38). HIV infection is also associated to pancytopenia and bone marrow failure (39, 40).

Whereas in VL patients the number of white and red blood cells and platelets is at least partially restored over time, this was not the case in the VL/HIV coinfected cohort. Both pathogens can infect hematopoietic stem/progenitor cells (HSPCs) and, at least in HIV infection, this can interfere with haematopoiesis (41). It is possible that the combined impact of *L. donovani* and HIV, via direct infection of HSPCs, or indirectly, via mechanisms such as high levels of chronic inflammation, becomes too detrimental for an adequate bone marrow output.

In summary, our clinical data suggest that better management of malnutrition and antiretroviral therapy could improve immune responses of VL/HIV patients and result in a more efficient control of parasite replication.

In addition to the follow-up of the clinical natural history of VL and VL/HIV infections, we also measured immunological markers to identify: a) mechanisms resulting in VL relapse; b) predictors of relapse. Since VL occurs in remote areas on Ethiopia, we focussed on markers that can be tested in a primary hospital setting.

Our results using a WBA to measure the production of antigen-specific IFNγ show that in contrast to those of VL patients, WBC from VL/HIV patients remain hyporesponsive as they fail to restore their capacity to produce IFNγ over time. The levels of IFNγ produced by WBC from VL/HIV patients without relapse after treatment was significantly higher as compared to those with relapse; however, these values were similar to those of VL patients at ToD, a time when these patients cannot control parasite replication. In the WBA, CD4^+^ T cells are the main source of IFNγ and this IFNγ has been shown to contribute to parasite killing (42, 43); it is therefore tempting to speculate that the inability of CD4^+^ T cells in VL/HIV patients to produce antigen-specific IFNγ over time plays a major role in the absence of efficient control of parasite replication after treatment. To identify the mechanisms resulting in suboptimal IFNγ, we tested the following hypotheses:

i. Low production of antigen-specific IFNγ is caused by low numbers of CD4^+^ T cells: our results show that VL/HIV patients fail to achieve normal CD4^+^ T cell counts. Since CD4^+^ T cells have been shown to be the main IFNγ-producing cells in the WBA (42, 43), we propose that the impaired antigen-specific production of IFNγ in VL/HIV patients is at least in part due to low CD4^+^ T cell counts. In contrast, CD4^+^ T cell counts were restored in VL patients after treatment. This shows that there is a normal recovery of CD4^+^ T cell counts in patients who are only infected with *Leishmania* parasites. It has been shown that despite ART, poor recovery of CD4^+^ T cell counts is common, especially in individuals who start ART with low CD4^+^ T cell counts (44–46); and results of a multi-country prospective cohort study in sub-Saharan Africa, showed a suboptimal recovery of CD4^+^ T cells despite sustained viral suppression on continuous first-line ART (47). In addition to bone marrow failure, reduced thymic output might also play a role in the low number of CD4^+^ T cells, and indeed, a recent study showed a clear correlation between decreased numbers of recent thymic emigrants and poor CD4+ T cell recovery in HIV patients (48). The study by Silva-Freitas et al. also suggested that in VL/HIV co-infected patients who do not relapse, more new emigrant T cells can be detected that might contribute to the control of parasite replication (49). In addition to its potential contribution to the low levels of IFNγ produced in the WBA, a poor CD4^+^ T cell recovery is also associated with persistent immune activation and inflammation (50); that in turn contribute to higher risks of several morbidities, as well as death (51, 52). Whereas to date, there are no effective alternative ART treatments that have been shown to increase the restoration of CD4^+^ T cell counts, it has been shown that early initiation of ART might boost the restoration of CD4^+^ T cell counts; furthermore inclusion of dolutegravir in first line of treatment could improve further CD4^+^ T cell recovery (53) through more rapid and sustained suppression of HIV replication. It also has a higher barrier to resistance making treatment failure less likely.
ii. Persistent inflammation contributes to T cell dysfunction and the subsequent low production of IFNγ (26). It has been well established that persistent inflammation remains in HIV patients despite viral suppression and increase in CD4^+^ T cell counts (54–56). *Leishmania* infections alone are also associated with persistent inflammation the current study shows that levels of cytokines such as TNF*α*, IL-6 and IL-8 remained increased 6-12 months post the end of anti-leishmanial treatment. In our cohort of VL/HIV patients, the levels were even further increased. It is therefore likely that this persistent inflammation contributes to T cell dysfunction.
iii. Low production of IFN*γ* is due to T cell exhaustion. In addition to inflammation, persistent antigenic stimulation results in T cell exhaustion, characterised by a progressive loss of efficient effector functions and co-expression of inhibitory receptors (26, 57, 58). Here, the immune check-point molecule PD1 was clearly expressed on CD4^+^ T cells from both patient cohorts at diagnosis. PD1 levels fell to be similar to those of controls in VL patients, accompanied by increasing levels of IFNγ secretion by whole blood cells, but remained high throughout follow-up in VL/HIV coinfected patients with low levels of antigen-specific IFNγ produced by whole blood cells. CD4^+^ T cells in VL/HIV patients thus continue to have hallmarks of exhausted cells: they express PD1 and produce low levels of IFNγ. Exhausted T cells do not lose all their effector functions, they become hypofunctional and can still proliferate and produce effector molecules (26, 57, 58). Whereas exhaustion is mainly seen as a dysfunctional state, exhausted T cells have been shown to at least partially contain chronic infections and play an important role in limiting immunopathology (26, 57, 58). It is therefore possible that during VL infections, activated T cells progressively become exhausted and that while still contributing to the control of parasite replication, they also contribute to prevent tissue damage. After successful treatment, with the clearance of the large majority of parasites and a progressively reduced inflammation, our results show that the frequency of these exhausted cells is restored back to control levels. In contrast, in VL/HIV patients, the persistent high antigenic stimulation and inflammation contribute to the maintenance of exhausted T cells.

We propose that in VL patients at ToD, low CD4^+^ T cell counts and T cell exhaustion play key roles in the impaired production of IFNγ, but that this is reversed at the end of successful anti-leishmanial treatment. In contrast, in VL/HIV patients, CD4^+^ T cell counts remain low and exhausted after anti-leishmanial treatment, resulting in severely impaired production of IFNγ that contributes to uncontrolled parasite replication and frequent relapses seen in VL/HIV patients.

Our study has identified 3 markers that were measured in a primary hospital in Ethiopia that are associated with VL relapse: low CD4^+^ T cell counts, low IFN*γ* production in a WBA, and high PD1 expression levels on CD4^+^ T cells. The multinomial logistic regression analyses show a good differentiation between all three groups (relapse at 3m, relapse at 6-12m and no relapse), and perhaps most clinically translatable is the clear distinction seen between the 3m and 6-12m groups. Importantly, these 3 measurements combine non-additively in the best prediction model (relapse at 3m), suggesting that these factors all interact in driving relapse in VL/HIV patients. These results provide a promising indication of the potential of widely available predictors from blood immunology tests to identify VL/HIV co-infected individuals most at risk of early relapse and need to be further validated in external cohorts of patients to ensure reproducibility. The simplicity of the prediction model will also lend itself readily to translation into a formula that can be calculated on site. This is extremely promising for such a setting, where individuals who are most at risk of relapse within 3 months could be identified and treated further: for these patients, in addition to an optimisation of their ART treatment and an improvement of their nutritional status, a longer anti-leishmanial treatment might be beneficial. Furthermore, the impaired production of IFNγ and high expression of PD1 suggest that immune therapy, through IFNγ administration (59) and/or PD1/PDL-1 blockade, might improve parasite killing and disease control in these patients.

## METHODS

### Patient recruitment

For this cross-sectional study, 25 healthy male non-endemic controls (median age 28.0±1.3 years old) were recruited among the staff of the University of Gondar, Ethiopia; and 99 male patients with visceral leishmaniasis (VL patients) were recruited from the Leishmaniasis Treatment and Research Centre (LRTC), University of Gondar. Forty-nine were HIV infected (VL/HIV) (median age 33.5±1.0 years old) and 50 were HIV uninfected (VL)(median age 25±0.8 years old). Patients age <18 years were excluded. The diagnosis of VL was based on positive serology (rK39) and the presence of *Leishmania* amastigotes in spleen or bone marrow aspirates (24). The diagnosis of HIV was in accordance with the Ethiopian national HIV screening test guideline. 46 VL/HIV patients were on ART at the time of VL diagnosis, the remaining three started ART at the end of the anti-leishmanial treatment. All treatments were administered according to the Guideline for Diagnosis and Prevention of Leishmaniasis in Ethiopia (22). At EoT, all VL patients were clinically cured, defined as follows: patients look improved, are afebrile, usually have a smaller spleen size and have an improved haematological profile. A test of cure (TOC) was used for VL/HIV patients to decide if they could be discharged from hospital; a negative TOC is defined as follows: patients look improved, afebrile, and usually have a smaller spleen size, parasitological cure (absence of amastigotes in splenic aspirates) and an improved haematological profile. When the TOC was still positive, treatment was continued until TOC becomes negative (22).

ART was provided according to the National guideline for comprehensive HIV prevention, care and treatment (60).

VL and VL/HIV patients were recruited at four different time points: time of diagnosis (ToD); end of treatment (EoT); 3 months post the end of treatment (3m); and 6-12 months post the end of treatment (6-12m). At the end of this 3-year study, all VL/HIV patients who had not relapsed during the 12 months follow-up were contacted by phone to find out if they had had any further relapse.

### Sample collection and processing

13ml of blood were collected by venepuncture and distributed as follows:

### 2.5ml in PAXgene tubes for RNA sequencing

8 ml in heparinised tubes: 3ml for the whole blood assay (WBA), 5ml to purify PBMC (18) (flowcytometry) and plasma (cytokines)

2.5ml in EDTA tubes (whole blood for: CD4^+^ T cell, white and red blood cell and platelet counts and plasma for viral load)

- WBA: soluble leishmania antigen (SLA) was prepared as described in (18).

- Flowcytometry: the following antibodies were used: CD4^FITC^, CTLA-4^APC^ and PD1^PE^ (eBioscience) (18). The data on PD1 expression are shown as the integrated Median Fluorescent Intensity (iMFI), which reflects the total functional response (61) more accurately than the frequency of expression.

- CD4^+^ T cell counts: 100µl of EDTA whole blood was stained with CD4^FITC^ and CD3^PerCP-eFluor® 710^ (eBioscience) for 15min at 4°C; red blood cells were lysed using BD FACS™ Lysing Solution for 5min at room temperature.

Acquisition was performed using BD Accuri™ C6 flow cytometry and data were analysed using BD Accuri C6 analysis software.

- IFN*γ* and IL-10 levels were measured in the supernatant of the WBA using IFN gamma and IL-10 Human ELISA Kit (Invitrogen) according to the manufacturer’s instructions.

- Inflammatory markers: Plasma was collected after gradient centrifugation of 5ml of heparinised blood as described in (18) and IFNγ, IL1β, IL2, IL4, IL6, IL8, IL10, IL12p70, IL13, TNFα and CRP plasma levels were measured by multiplex assay using U-PLEX Proinflam Combo 1, V-PLEX Proinflammatory Panel1 (62) Kit and V-PLEX Human CRP following the manufacturer’s instructions (Meso Scale Diagnostics, USA).

- HIV viral load: Plasma was isolated by centrifuging 2ml of EDTA whole blood and frozen at −80°C. HIV viral load was measured in the Central Laboratory of the Amhara Public Health Institute, Bahir Dar, by using Abbott RealTime HIV-1 Qualitative (m2000sp), according to the manufacturer’s instructions.

- White and red blood cell, and platelet counts were measured using a Sysmex XP-300T^M^ automated haematology analyser, (USA) following the manufacturer’s instruction.

- Liver (serum aspartate aminotransferase (SGOT) and serum glutamic pyruvic transaminase (SGPT)) and renal function (blood urea nitrogen (BUN) and creatinine) tests were performed as part of the routine management practice.

### Relapse rate

Log-rank tests for duration of follow-up at event end points provided two-sided *p*-values; Kaplan-Meier curves are presented for visual interpretation. The primary outcome survival until three years of follow-up was completed; relapse was the only censoring event. Censoring events were reported at the pre-planned follow-up period (3, 6-12 and after 3 years) at which they were identified. Cox proportional-hazards regression analysis was used to estimate hazard ratios and 95% confidence intervals

### mRNA

2.5 ml of blood was collected in PAXgene blood RNA tubes, RNA extracted using the PAXgene 96 blood RNA kit (Qiagen) and Globin mRNA depleted using the GLOBINclear kit (Ambion). Sequencing libraries were prepared using the KAPA Stranded mRNA-Seq Kit (Roche) with 10 PCR cycles, then sequenced as 75bp paired-end reads on the Illumina HiSeq 4000 platform. Sequencing reads were mapped with Salmon v.1.30 against concatenated sequence of human gencode transcriptome release 34, transcripts for *L. donovani* LV9 from TriTrypDB release 46 and transcript data for an Ethiopian HIV type 1C virus (Genbank Accession U46016)

- type 1C represents over 90% of HIV infections in Ethiopia. Pseudo-counts were imported into R v4.0.3 using the tximport v1.18.0 and transformed into lengthscaledTPM. Total HIV expression was quantified as the sum of counts across HIV transcripts, and total *Leishmania* expression was quantified as the sum across all LV9 transcripts with the exception of feature LdLV9.27.2.206410, an 18S rRNA gene to which human transcripts also map.

### Logistic regression

A multinomial logistic regression analysis was performed in R (version 4.0.3) (R Core Team, 2020), using raw CD4 counts, IFNγ and PD1 measured at EoT to predict relapse time for individuals. After removing missing data, a cohort of 34 individuals were allocated to three potential categories based on time (in months) to relapse after EoT: “Relapse at 3m” (n=10), “Relapse at 6-12m”, (n=7) and “No Relapse in Study” (n=17). Performance of the model of best fit was assessed by plotting a receiver operating curve (ROC) for predicting relapse at 3m and 6-12m. Area under the ROC curve (AUC) was then calculated with a 95% confidence interval (CI) for both groups.

### Statistical analysis

Data were evaluated for statistical differences as specified in the legend of each figure. The following tests were used: one-way ANOVA, Mann-Whitney, Kruskal-Wallis or Spearman’s rank test. Differences were considered statistically significant at *p*<0.05. Unless otherwise specified, results are expressed as median±SEM.

*=p<0.05, **=p<0.01, ***=p<0.001 and ****=p<0.0001.

### Study approval

This study was approved by the Institutional Review Board of the University of Gondar (IRB, reference O/V/P/RCS/05/1572/2017), the National Research Ethics Review Committee (NRERC, reference 310/130/2018) and Imperial College Research Ethics Committee (ICREC 17SM480). Informed written consent was obtained from each patient and control.

## AUTHOR CONTRIBUTIONS

All authors discussed the results and contributed to the final manuscript. Study conception and design: PK, YT, IM; Acquisition of data: YT, EA, TM, PK; Analysis and interpretation of data: YT, TM, EA, CJS, SF, RW, MK, GPT, ML, IM, JAC, PK; Drafting of manuscript: PK, YT, JAC, IM

## ACKNOWLEDGMENTS

We are grateful to the staff of the Leishmaniasis Research and Treatment Centre, University of Gondar, for their support. We also would like to thank Mandy Sanders and Siobhan Austin-Guest at the Wellcome Sanger Institute for supporting and co-ordinating sequencing work; and Profs. CRM Bangham, Imperial College London, EM Riley, University of Edinburgh, and M. Berriman, Wellcome Sanger Institute, for helpful discussions. YT is funded by a Wellcome Trust Training Fellowship in Public Health and Tropical Medicine (204797/Z/16/Z). JAC is funded by Wellcome via core funding of the Wellcome Sanger Institute (grant 206194). MK is funded by a Wellcome Trust Sir Henry Wellcome Fellowship (206508/Z/17/Z).

