## Supplementarydatatakeleetal. for "Immunological factors, but not clinical features, predict visceral leishmaniasis relapse in patients co-infected with HIV"

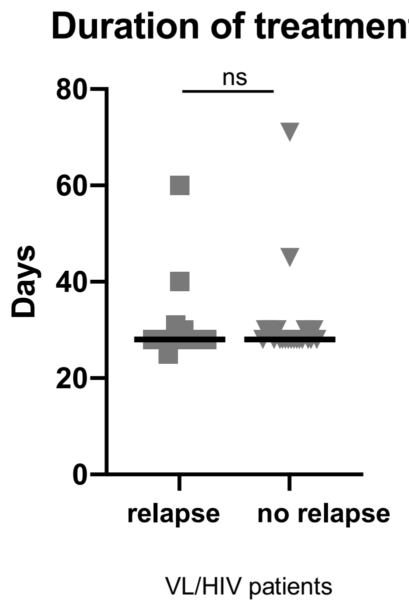


**Figure S1: Duration of treatment:** comparison of the duration of initial VL treatment (at ToD) between VL/HIV patients who relapsed and those who didn’t relapse during follow-up.

Each symbol represents the value for one individual, the straight lines represent the median. Statistical differences were determined using a Mann-Whitney test.

ns=not significant.

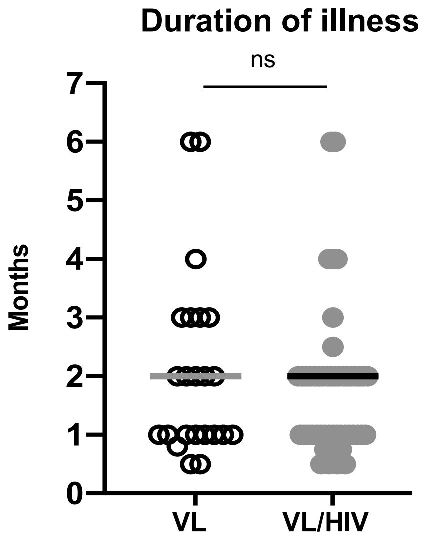


**Figure S2: Duration of illness:** comparison of the duration of illness as defined by the duration of time since the patients have started experiencing symptoms such as enlarged abdomen, fever, fatigue and loss of weight.
Each symbol represents the value for one individual, the straight lines represent the median. Statistical differences were determined using a Mann-Whitney test.
ns=not significant.

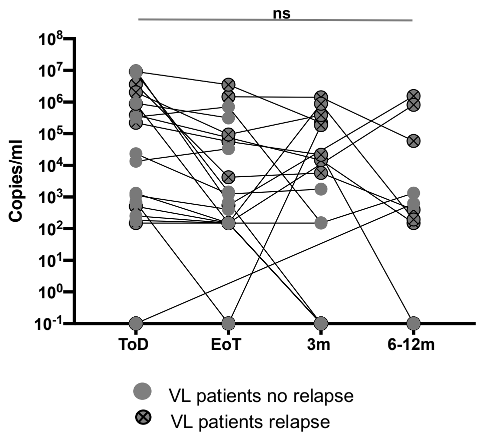


**Figure S3: viral load:** viral load (copies/ml) were measured longitudinally in the plasma of 35 VL/HIV patients who had at least 2 measurements of their viral load over the duration of the study.

Grey full circle=VL/HIV who did not relapse during the study. Grey full circle with a cross=VL/HIV who relapsed during the study.

Data are shown for patients who had at least 2 measurements during the course of the study.

Statistical differences were determined using a one-way ANOVA.

ns=not significant.

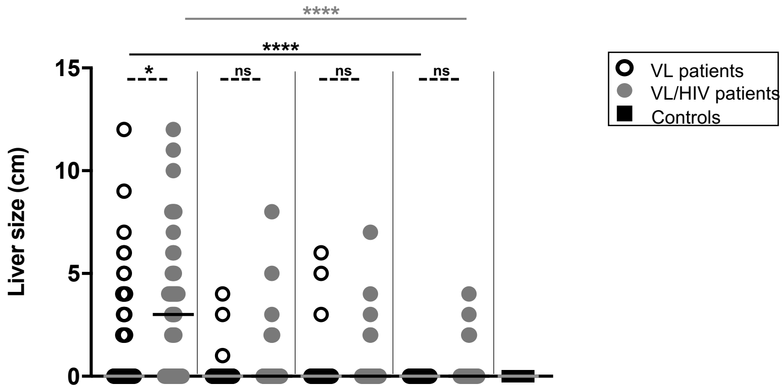


**Figure S4: liver size** was measured in cm below the costal margin on VL (ToD: n=50, EoT: n=45, 3m: n=36, 6-12m: n=26), VL/HIV (ToD: n=49, EoT: n=39, 3m: n=32, 6-12m: n=26) patients and controls (n=25) and controls (n=25).
Each symbol represents the value for one individual, the straight lines represent the median. Statistical differences between VL and VL/HIV patients at each time point or between no relapse and relapse were determined using a Mann-Whitney test and statistical differences between the 4 different time points for each cohort of patients were determined by Kruskal-Wallis test.
ToD=Time of Diagnosis; EoT=End of Treatment; 3m=3 months post EoT; 6-12m=6-12 months post EoT. ns=not significant.

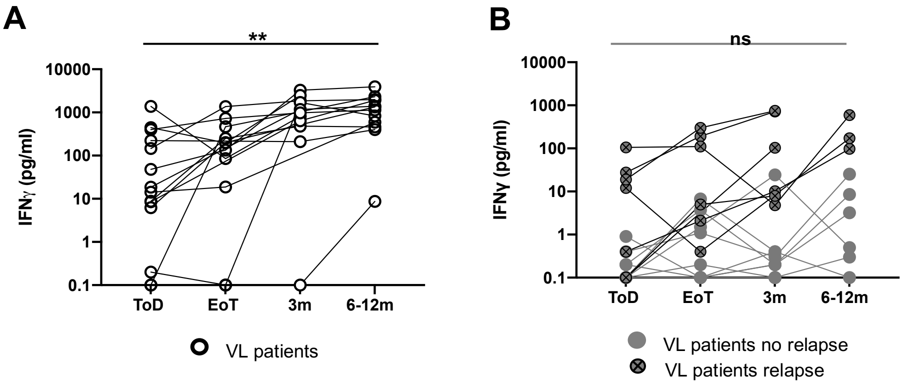


**Figure S5: Whole blood assay: longitudinal antigen-specific production of IFNγ:** Whole blood cells from **A.** VL patients (ToD: n=14, EoT: n=11, 3m: n=11, 6-12m: n=12) and **B.** VL/HIV patients (ToD: n=17, EoT: n=16, 3m: n=16, 6-12m: n=9) were cultured in the presence of SLA and IFNγ was measured by ELISA in the supernatant after 24hrs.

Grey full circle=VL/HIV who did not relapse during the study. Grey full circle with a cross=VL/HIV who relapsed during the study.

Data are shown for patients who had at least 2 measurements during the course of the study.

Each symbol represents the value for one individual.

Statistical differences were determined using a one-way ANOVA.

ToD=Time of Diagnosis; EoT=End of Treatment; 3m=3 months post EoT; 6-12m=3 months post EoT. ns=not significant.

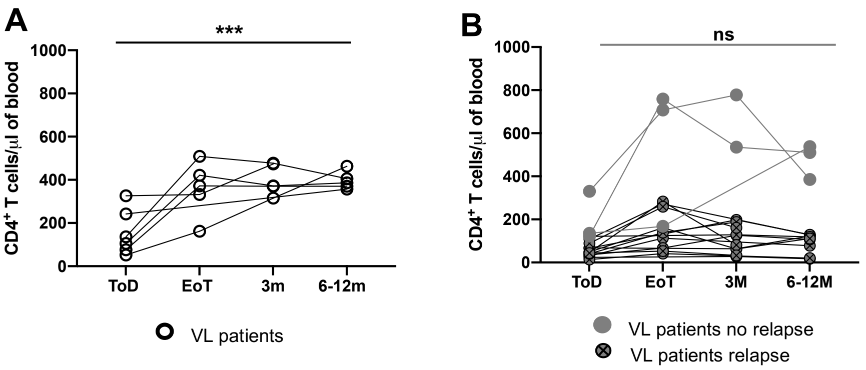


**Figure S6: Longitudinal CD4$^+$ T cell counts:** CD4$^+$ T cell counts were measured by flow cytometry in the blood of **A.** VL (ToD: n=6, EoT: n=5, 3m: n=5, 6-12m: n=5) and **B.** VL/HIV patients (ToD: n=16, EoT: n=12, 3m: n=13, 6-12m: n=12).
Grey full circle=VL/HIV who did not relapse during the study. Grey full circle with a cross=VL/HIV who relapsed during the study.
Data are shown for patients who had at least 2 measurements during the course of the study.
Each symbol represents the value for one individual. Statistical differences were determined using a one-way ANOVA.
ToD=Time of Diagnosis; EoT=End of Treatment; 3m=3 months post EoT; 6-12m=6-12 months post EoT. ns=not significant.

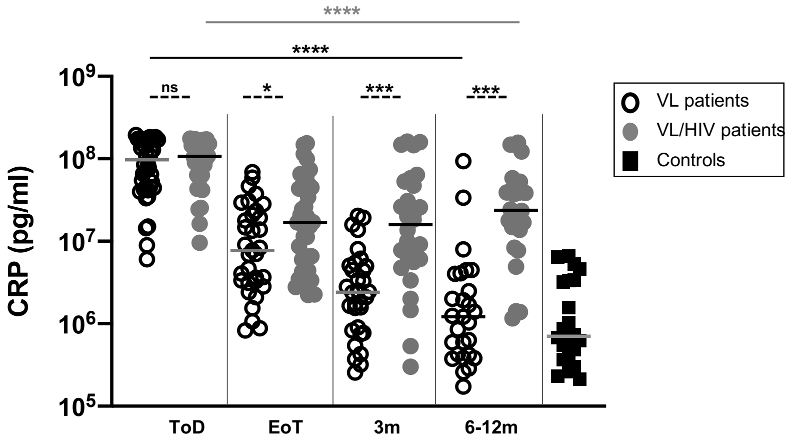


**Figure S7: CRP levels in the plasma of VL and VL/HIV patients:** CRP levels were measured in the plasma isolated from the blood of VL (ToD: n=42, EoT: n=33, 3m: n=32, 6-12m: n=27), VL/HIV patients (ToD: n=29, EoT: n=39, 3m: n=28, 6-12m: n=28) and controls (n=22) by multiplex assay.
Each symbol represents the value for one individual, the straight lines represent the median. Statistical differences between VL and VL/HIV patients at each time point or between no relapse and relapse were determined using a Mann-Whitney test and statistical differences between the 4 different time points for each cohort of patients were determined by Kruskal-Wallis test.
ToD=Time of Diagnosis; EoT=End of Treatment; 3m=3 months post EoT; 6-12m=6-12 months post EoT. ns=not significant.

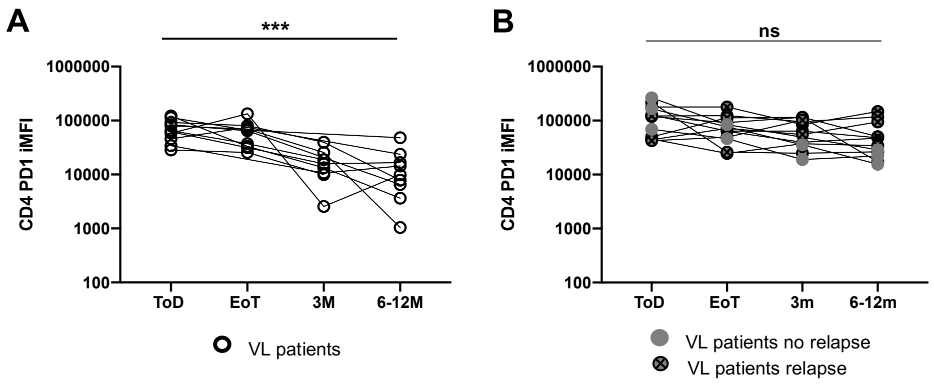


**Figure S8: Longitudinal follow-up CD4 PD1 iMFI:** CD4 PD1 iMFI was measured by flow cytometry in the PBMCs of **A.** VL (ToD: n=11, EoT: n=10, 3m: n=8, 6-12m: n=9) and **B.** VL/HIV patients (ToD: n=11, EoT: n=12, 3m: n=11, 6-12m: n=11). Grey full circle=VL/HIV who did not relapse during the study. Grey full circle with a cross=VL/HIV who relapsed during the study.

Data are shown for patients who had at least 2 measurements during the course of the study.

statistical differences were determined by Kruskal-Wallis test.

ToD=Time of Diagnosis; EoT=End of Treatment; 3m=3 months post EoT; 6-12m=6-12 months post EoT. ns=not significant.
